# The Roles of Neighborhood Composition and Autism Prevalence on Vaccination Exemption Pockets: A Population-wide Study

**DOI:** 10.1101/323451

**Authors:** Ashley Gromis, Kayuet Liu

## Abstract

The number of children entering schools without mandated vaccinations has increased in high-income countries due to the rise of nonmedical exemptions from school vaccination requirements. Herd immunity is threatened when unvaccinated children are concentrated in spatial pockets. It is often assumed that these exemption clusters are merely the result of population composition. On the other hand, despite the role of vaccine-autism controversy to the current wave of anti-vaccine movement, we do not know if exemption clusters are associated with local autism rates. Our spatial analysis of California shows that while racial/ethnic composition is associated with the locations of large exemption pockets, other sociodemographic factors and access to health care resources have limited geographical span. We decouple the race/ethnicity effect from that of unobserved socioeconomic status by examining families in poverty. Using unique address-level data on the locations of the majority of children with an autism diagnosis, we show that the prevalence of autism is not associated with the locations of large pockets of vaccination exemptions. In addition, we find charter schools in most exemption clusters; potential spillovers from charter schools to neighboring public schools are evaluated. Exemption pockets are not merely the result of population composition and community-level interventions are needed to maintain herd immunity.

**Highlights:**

- Autism prevalence rates are not associated with the locations of large exemption pockets.
- The average exemption rate in charter schools (7.5%) was higher than private schools.
- Proportion non-Hispanic white has the strongest association with large exemption clusters.
- Population composition cannot fully explain the exemption clusters.

## INTRODUCTION

Parental concerns over vaccine safety are suggested to have overtaken health care access as the primary obstacle to immunization in high-income countries (1). Anti-vaccine beliefs have been shown to be difficult to change, and policy changes to tighten immunization requirements are often met with resistance (2, 3). Concentrations of exempted children in certain areas increase the risk of vaccine-preventable disease outbreaks (4, 5). When other interventions are ineffective or infeasible, preventing exemption can preserve herd immunity.

This study seeks to identify mechanisms underlying vaccination exemption clusters. First, we explore how the discredited association between vaccines and autism might have affected the spatial patterns of exemptions. Despite the importance of the vaccine-autism controversy to the current wave of anti-vaccine movement, this is the first study to use address-level data of autism prevalence to study its relationship with exemption clusters. Like vaccination exemptions, autism clusters spatially (6, 7) and have a positive socioeconomic (SES) gradient (8, 9). While many studies demonstrated that vaccines do not cause autism (see 10), a substantial number of parents are still concerned about the alleged link (11, 12). Research on vicarious learning (13) suggests that local autism prevalence rates may have a positive association with exemption rates.

Secondly, previous studies have shown that exemptions are more prevalent in predominately white, high SES neighborhoods (14–21). This study asks a different set of questions: is the sorting of children by SES into certain areas sufficient to explain large pockets of exemptions that span multiple schools? If so, why do neighborhoods with residents of similar SES have drastically different exemption rates? Moreover, do some socio-demographic factors have a limited geographical span while others are more important for large clusters? Understanding the geographical span of the sociodemographic correlates of exemptions will shed light on the underlying mechanisms generating these clusters and has policy implications.

## METHODS

### Data

*Personal Beliefs exemptions*. This study uses the California Department of Public Health (CDPH) annual data on non-medical, personal belief exemptions (PBEs) from school vaccination requirements from 1998-2014.^1^ Kindergartens and licensed childcare centers with enrollments of more than10 students are required to report exemptions to the CDPH. In 2014, a physician’s signature was required to file a PBE. This resulted in a drop in PBEs reported to the CDPH, including those that might have been filed out of convenience. Our spatial analyses focus on 2014 to yield a conservative estimate of exemptions that were primarily driven by vaccine-related beliefs.

*Autism*. We use data from California because it is the only U.S. state that uses a centralized system of both the assessment of and service provision to over 70% of individuals with autism (23). Information from the Department of Developmental Disorders on the addresses of children with autism from 1992-2011 provides a unique opportunity to study the associations between autism prevalence and vaccination exemptions.^2^ Further details of the data are described elsewhere (8). We count the number of children under age 10 with an autism diagnosis within a 500-child^3^ radius around a school or childcare center. The flexible geographical unit allows us to compare rural and urban areas. To compare the effect of autism incidence with autism advocacy efforts, county-level counts of autism advocacy organizations are collected from Internal Revenue Office’s records of tax-exempt organizations.

*Pertussis*. Outbreaks of vaccine-preventable diseases are expected to have the opposite effect of autism. Immunization rates have been shown to increase after epidemics (25). Rates per 1000 population of pertussis in California’s counties from 2000 to 2010 are obtained from CDPH’s Immunization Branch.

*Proximity to physicians*. Access to health care is traditionally a major determinant of immunization level. Access to physicians is measured by the count of general, family, and pediatric physicians within a school’s 500-child radius. These counts are created from physician data provided by the American Medical Association.

*Other selection processes*. Selection into private schools and charter schools^4^ with alternative pedagogical orientations may increase PBE rates (21, 26). Other local factors such as the availability of alternative medicine practitioners have been suggested to increase exemptions (27–29). We construct the counts of chiropractors and acupuncturists within a school’s 500-child radius based on licensing records from the California Department of Consumer Affairs.

*Socioeconomic factors*. Various mechanisms could account for positive SES gradient of PBEs, such the perceived risks of contagious diseases (28). Despite the many findings on SES and PBEs, the relationship between SES and vaccine-related *beliefs* is unclear (30). Low SES and ethnic minority parents have more vaccine safety concerns than high SES parents, even though they still vaccinate their children (11, 31). That said, a concentration of high SES individuals among the small number of parents who actually seek PBEs is sufficient to generate the positive SES gradient.

We measure SES by the average maternal education level and property values within the school’s 500-child radius described above. Education is measured as the highest year of schooling completed by the mother at the time of birth. This information is obtained from address-level data in the California Birth Statistical Master File between 1992 and 2007. Median property values for the full study period are interpolated using the Censuses’ block-group level data and logged. Racial composition is represented as the percentage non-Hispanic white residents within the 500-child radius, using interpolated Census block-group level data. Unfortunately, other racial/ethnic categories cannot be considered as their small numbers in certain areas would have yielded unstable estimates. California’s Department of Education provides enrollment breakdowns by race/ethnicity for public schools and are used to calculate white isolation indices (32) at the school district level.^5^ These indices measure the extent to which non-Hispanic white students are exposed only to one another. Population density is interpolated using the Censuses’ block group level data and logged.

### Analysis

We conduct our analysis in two stages. First, we use negative binomial regression with random effects to yield baseline estimates of covariates’ effects on the annual counts of PBEs at the school-level.^6^ All covariates are lagged 3 and 6 years for childcare centers and kindergartens, respectively, to account for the majority of vaccination decisions taking place before a child enters school.^7^

After obtaining the school-level estimates described above, our main analysis uses Kulldorff Spatial Scan Statistic (33) to map clusters of high PBE rates among public schools,^8^ adjusting for the spatial distributions of the covariates. Maximum likelihood estimation identifies circular areas that encompass schools with an increased probability of PBEs compared to the state-wide PBE rate. Certainly, not all schools inside the clusters have high PBE rates. However, this method systematically identifies areas in which all children have a heightened risk of exposure to non-vaccinated children (34).

## RESULTS

### School-Level Correlates

Table 1 shows the results of our school-level model. Count of autism diagnoses has the expected positive effect on PBEs at the school level. Count of acupuncturists is statistically insignificant once the other covariates are included in the model. PBE rates increase with physician density, which is consistent with a previous study (28).^9^ Maternal education and percentage non-Hispanic white have the expected positive associations with PBEs. Property values are not associated with PBEs once the effects of other covariates are controlled for.

**Table 1.** **Correlates of PBEs among public school kindergartens, 1998-2014^1^**

|  | Year-adjusted<sup>2</sup><br>IRR<sup>3</sup> | 95% C.I. |  | Fully-adjusted<br>IRR | 95% C.I. |  |
| --- | --- | --- | --- | --- | --- | --- |
| Autism diagnoses | 1.013*** | 1.007 | 1.019 | 1.006*** | 1.000 | 1.012 |
| Acupuncturists<sup>4</sup> | 1.038*** | 1.030 | 1.045 | 0.996*** | 0.988 | 1.003 |
| Chiropractors<sup>4</sup> | 1.043*** | 1.038 | 1.048 | 1.029*** | 1.023 | 1.034 |
| Physicians | 1.010*** | 1.008 | 1.012 | 1.004*** | 1.002 | 1.006 |
| Mother's education | 1.532*** | 1.505 | 1.560 | 1.190*** | 1.163 | 1.218 |
| Property values (ln) | 1.651*** | 1.575 | 1.730 | 0.974*** | 0.926 | 1.025 |
| % white<sup>5</sup> | 1.339*** | 1.329 | 1.348 | 1.217*** | 1.205 | 1.230 |
| Pop. density (ln) | 0.683*** | 0.669 | 0.698 | 0.782*** | 0.766 | 0.799 |

Notes:
<sup>1</sup> 2-level random intercept negative binomial regression model. N=89,550 school-years. Year indicator variables included in all models but not shown. All independent variables lagged 6 years. We estimated spatial lag models as robustness checks in light of potential spatial auto-correlation in the errors. Estimates and statistical significance of covariate effects do not differ substantially. We choose not include spatial lags in here because our aim is to map rather than statistically control for the unexplained spatial clustering of PBEs.
<sup>2</sup> Adjusted for year indicator variables.
<sup>3</sup> Incident Rate Ratio
<sup>4</sup> To ensure that proximity to alternative healthcare practitioners is not due to a spurious effect of proximity to business areas, we also construct measures of the densities of randomly selected licensed professionals, as well as the density of veterinarians, which should not be associated with beliefs about vaccination. Proximity to randomly selected licensed professionals did not have a statistically significant effect. Count of veterinarians has a significant, positive effect on PBEs, net of other variables (IRR=1.02, p<0.001). We thus cannot rule out that unmeasured neighborhood-level characteristics could also explain the positive associations between chiropractors and PBE rates.
<sup>5</sup> Measured in 10% increments (deciles) for readability
\*\*\*p < .001

### PBE Pockets

We now turn to our main questions: are the large PBE pockets spanning multiple schools the product of the spatial distributions of socio-demographic factors? Are local autism rates associated with the locations of the large PBE clusters?

Herd immunity requires immunization levels of about 90%—higher for extremely infectious diseases, e.g., pertussis, and slightly lower for others (35). In 2014, the average PBE rate among kindergarteners in California was still far below 10%.

Yet the concentration of PBEs in a small number of schools is the issue. Private schools, particularly those adopt alternative education methods, have been shown to have higher rates of PBEs than public schools (26, 36). The average PBE rates in California’s private schools and non-charter public schools in 2014 were 5.2% and 2.1%, respectively. The average PBE rate of charter schools was as high as 7.5%, even higher than private schools. The Lorenz curves in Figure 2 show that, within each school type, schools comprising of 20% of students contribute to as much as 70% of PBEs.

**Figure 1.**
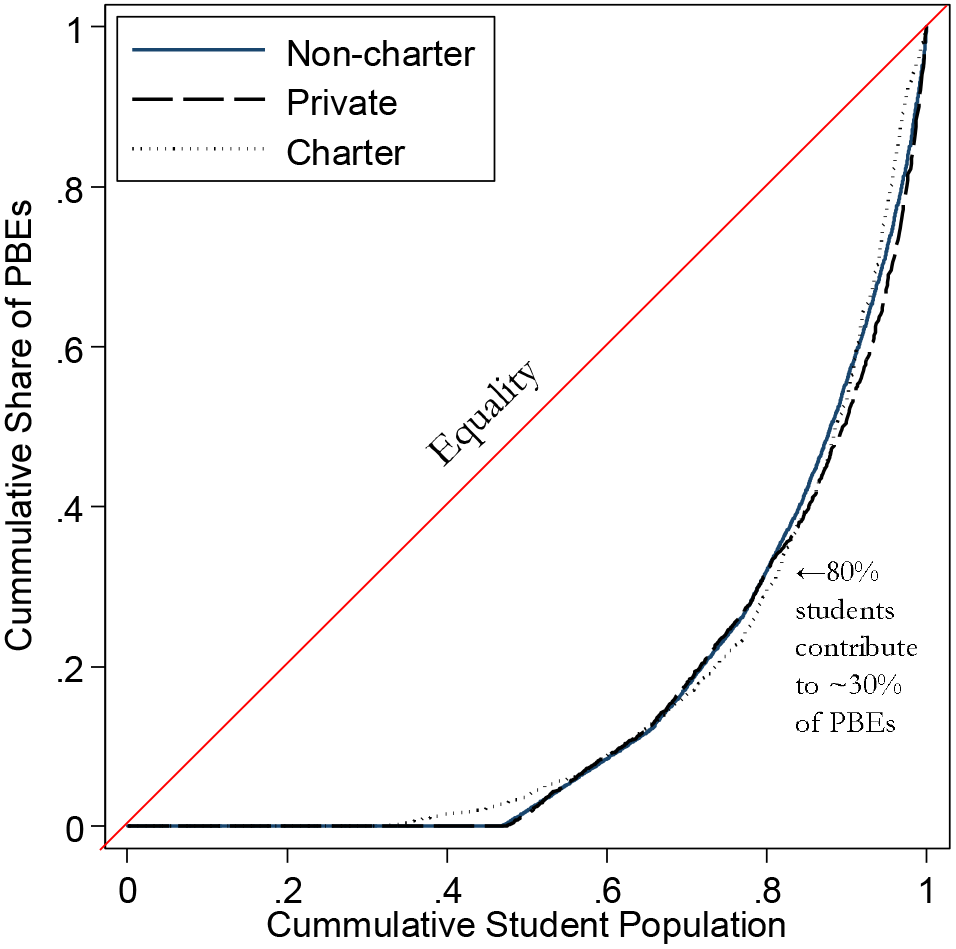
Lorenz curves of PBE distributions in 2014 by School Type.

The shaded circular regions in Figure 1A show where the high-risk areas are located without adjusting for any covariates: they have statistically significantly higher relative risk of PBEs than the non-shaded areas. These clusters of high PBE rates among public schools were not confined in certain areas but distributed all across the state, from the sparsely populated area in the north, to areas in the Central Valley, to coastal urban centers (e.g., San Francisco, Santa Clara, Santa Barbara, Los Angeles, and San Diego).

**Figure 2.**
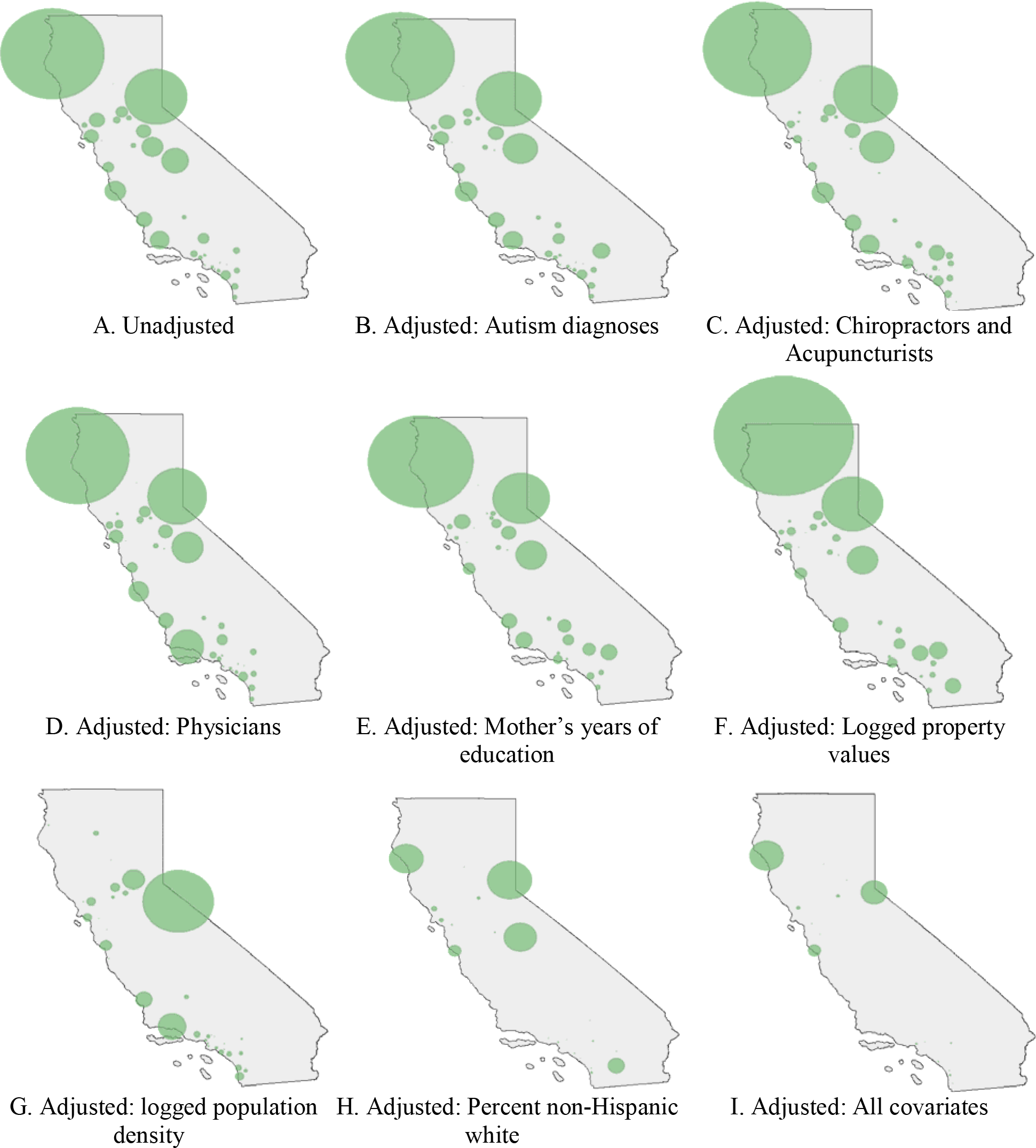
Unadjusted and adjusted spatial clusters of PBEs across public schools, 2014.

Figures 1B-H show the spatial clustering of PBEs across schools after adjusting for each of the school-level covariates. If the spatial clustering of PBEs is completely driven by a covariate’s spatial distribution, the adjustment will eliminate the PBE clusters from the corresponding figure.

*Autism and Pertussis*. Adjusting for spatial distribution of children with autism has almost no effect on the PBE pockets spanning multiple schools (Figure 1B), even though proximity to children with autism has a statistically significant effect in the school-level model. The major spatial cluster of autism was located in Los Angeles (6). In contrast, high PBE rates are found in other cities across California.

It is possible that the saliency of autism is not about knowing a child with autism but its advocacy. Our alternative specification uses count of autism awareness organizations. Because the count of organizations is at the county level, we used a three level mixed-effects Poisson model. The result shows that the locations of autism advocacy organizations have no statistically significant effect on PBEs (Table A2 in Appendix).

The same three-level mixed-effects model also shows that the association between pertussis outbreaks at the county level and PBEs is in the expected direction. An additional case of pertussis per 1000 among children aged 0-6 decreases PBE rates by 3.8%, net of other factors. However, the effect is only marginally significant statistically (p=0.086).

*Proximity to health care practitioners*. Adjusting for the locations of chiropractors and acupuncturists has almost no effect on the large pockets of PBEs (Figure 1C). Adjusting for counts of physicians similarly has negligible effect (Figure 1D).

*Socio-demographic factors*. Figure 1E shows that controlling for maternal education has a surprisingly small impact on the multiple-school clusters despite its strong association with PBE rates in the school-level model. In contrast, Figure 1F shows that while logged property values are not associated with PBEs at the school level, adjusting for their spatial distribution reduces the size of the clusters in the coastal urban areas. Controlling for logged population density shrinks some of the clusters, especially the cluster in the north (Figure 1G).

Among the covariates, adjusting for the percentage of non-Hispanic white has the most substantial impact in reducing both the number and sizes of PBE clusters (Figure 1H). This strong percent white effect is consistently observed across years (Figure A1 in Appendix) and therefore cannot be explained by idiosyncrasies of the 2014 data.

*Exploring the percent white effect:*. Why does percent non-Hispanic white have such a strong impact while mother’s years of education, which has a substantial effect size in the school level model, has almost no impact on reducing multiple-school clusters? In theory, any PBE-related spatial processes spanning multiple schools *and* correlated with percent non-Hispanic white could be responsible. The diffusion of vaccine skepticism over racially homophilous (i.e., similar) social networks is an example. We later discuss why there are reasons to expect spatially embedded, dense social networks are particularly relevant to vaccine refusals. While our data do not allow us to directly test the social diffusion of vaccine skepticism, the following robustness checks attempt to address some of the alternative explanations.

First, we examine the effect of percent white in a homogeneous population with regard to SES—children in Head Start programs. Percent non-Hispanic white could be associated with unmeasured SES. To be eligible for Head Start, families generally must earn less than or equal to the federal poverty line. If the percent non-Hispanic white result in Figure 1H is merely picking up unmeasured SES effects, we should expect it to have no effect on PBE rates among Head Starts.

However, Figure 3A shows the opposite result— among all school types, percentage non-Hispanic white has the *strongest* effect on Head Starts’ PBE rates. A 10% increase in percent non-Hispanic white in the area around a Head Start is associated with on average a 37% increase in PBEs, controlling for the effects of other covariates.

**Figure 3.**
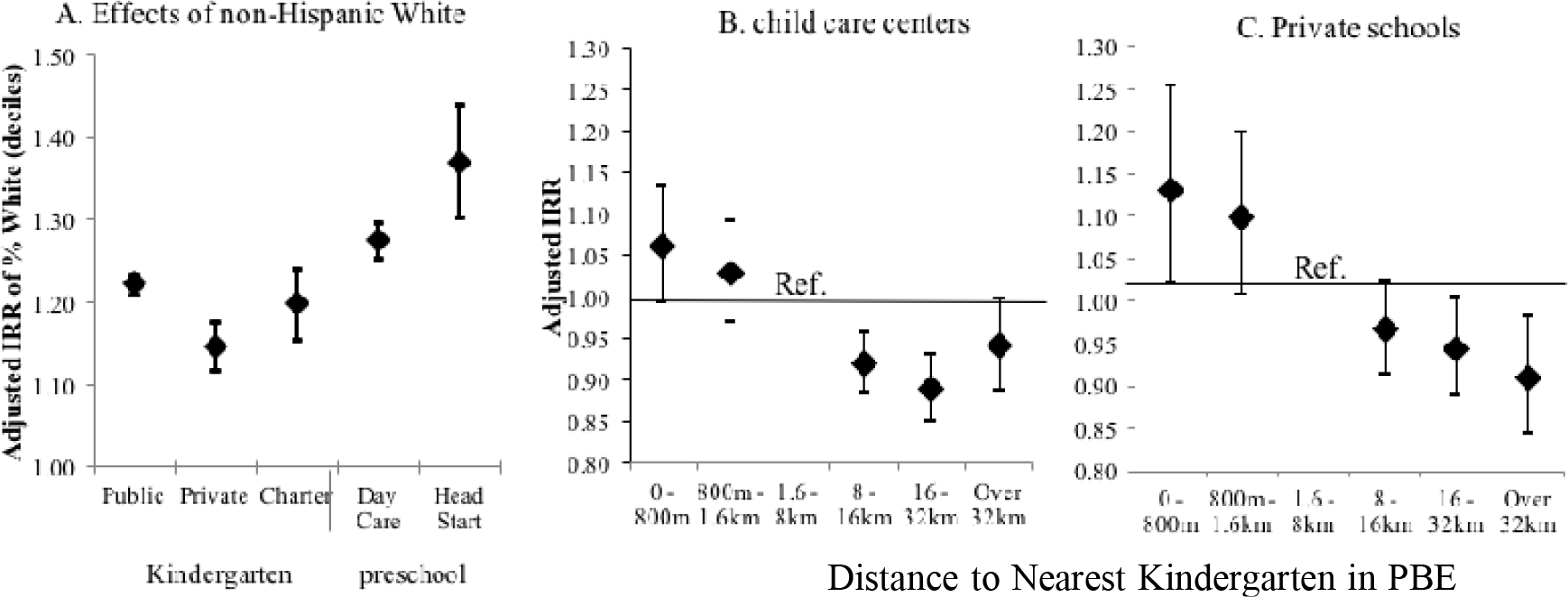
Ruling out alternative explanations of the association between percent non-Hispanic white and PBE clusters across schools. (A) Effect of percent non Hispanic white by school type; effects of distance to nearest public kindergarten in PBE cluster on PBE rates in (B) child care centers and (C) private schools. Note: Incidence Risk ratios (IRRs) estimated via negative binomial regression and adjusted for mother’s education, logged property values, logged population density, counts of physicians, chiropractors, acupuncturists, and autism diagnoses, and yearly indicator variables. Public schools in clusters only include those with statistically significantly higher relative risk of PBEs after adjusting for covariates. 1998-2014: Public schools N=84,303; private schools: N=27,378; charter schools: N=5,247; Head Start centers: N=6,049. 2010-2014: Childcare centers: N=36,850

Second, we ask whether the percent white effect on the clustering of PBEs could be explained by residual segregation patterns. If families are more segregated by race/ethnicity than education, it could explain why mother’s years of education only have localized effects. We calculated Moran’s I, a spatial correlation statistic, at the Census block-group level for both mother’s education and percent non-Hispanic white. According to Moran’s I, California was similarly segregated by being non-Hispanic white and by mother’s years of education (results available upon request). Hence, residential segregation alone cannot explain the substantial percent white effect on PBE pockets.

The above analysis does not mean that racial segregation does not matter. It just shows that segregation patterns alone do not explain why the effect of percent white can span large areas than that of mother’s years of education. In fact, Table A4 in the Appendix confirms that school districts with greater isolation of non-Hispanic white children (i.e., a high isolation index) have higher PBE rates than those districts that are more racially mixed.

Third, we test whether school-district factors may be responsible for the spatial association between non-Hispanic white population and PBEs. School district boundaries do not structure enrollment in child care centers and therefore should not influence their PBE rates. However, Figure 3B shows child care centers near kindergarten PBE clusters are more likely to have high PBE rates, controlling for the effects of socio-demographic variables. It is possible that this result was driven by unvaccinated kindergarteners having siblings that are also unvaccinated in nearby child care centers. It is less common, however, to have one sibling go to a public school while the other attends a private school. We repeated the same analysis with private schools and still find PBE rates of private schools to be statistically significantly higher if they are close to public kindergarten PBE clusters (Figure 3C).

*Charter Schools*. Figure 1I shows the effects of adjusting for all covariates. 48% of the PBE clusters still consist of multiple schools, although they now cover much smaller areas. Among the multi-school clusters, 87.5% include at least one charter school.

To test whether the spatial clusters are mainly driven by the charter schools, we removed them from the analysis and examined the PBE clusters only among non-charter public schools (Figure A2 in the Appendix). Adjusted for the spatial distributions of all covariates, PBEs continue to be unevenly distributed across non-charter public schools, with some clusters in the same locations as when charter schools are included.

We use our longitudinal data to test for potential spillovers from charter to non-charter schools: Table A5 in Appendix reports the effect of the opening of a charter school on PBE rates in nearby public schools. A fixed effects model shows that the public schools in PBE clusters have statistically significantly higher PBE rates the year after the opening of the charter school in the cluster than the year before.

## DISCUSSION

As of 2016, 47 U.S. states permit religious and/or philosophical exemptions (37). California now prohibits PBEs. However, disallowing PBEs seems to have driven some parents to medical exemptions (38). Banning PBEs could also concentrate unvaccinated children in independent study schools (39, 40). It is often assumed that spatial clustering of behaviors is merely the result of population composition or uneven health care access; our findings suggest that it is not the case for PBEs. Moreover, the effects of sociodemographic factors are shown to have different geographical span. The implication is that the spatial patterns of PBEs are likely the combined result of social processes operating at varying geographical scales. Interventions that go beyond the individual-level and address these social processes are needed to prevent exemption pockets.

Specifically, the seemingly unexplained increase in autism diagnoses (but see 8, 41) has fueled popularity of the belief that vaccines can cause autism. Not surprisingly, parents who file PBEs were concerned about vaccine safety (42, 43). Nonetheless, this study shows that there is little overlap between local autism prevalence and the locations of the large pockets of PBEs.

Instead, racial homogeneity, measured by percent non-Hispanic white, has a uniquely strong association with large PBE clusters while other socio-demographic factors show more local effects. We speculate that spatially embedded network interactions are behind this result. Social networks can substantially influence health decisions (44). In particular, dense social networks are crucial to the adoption of rare or risky behaviors (45), and are often racially homophilous (46). Dense networks tend to be spatially clustered and thus can exacerbate the clustering of risky behaviors (47). Such endogenous processes of social influence also explain why neighborhoods similar in SES can have drastically different PBE rates. Anti-vaccine attitudes are notoriously difficult to change (48) but network interventions (49) targeting communities may be able to change norms and prevent exemption pockets. Lastly, the charter school movement likely had an unintended consequence of further concentrating children with vaccine exemptions. Future debates about school choice should also consider its consequences for public health.

This study is limited by the lack of individual-level data. Although robustness checks were used to evaluate some alternative explanations of our results on race/ethnicity, detailed network data are required to directly examine the diffusion of vaccine refusals and the role of race/ethnicity. While lacking individual-level details, the statewide data allow us to isolate the spatial correlates of vaccination exemption at different geographical scales. Future research combining epidemiological and individual level data can shed light on the specific mechanisms.

## Footnotes

1 Although PBE status is indicative of lack of immunization or under immunization, our data do not tell us how many required vaccinations a child is lacking. See (22) for coverage estimates.

2 The DDS does not provide services to people with Asperger Syndrome or Pervasive Developmental Disorder-Not Otherwise Specified (PDD-NOS).

3 The radius is calculated based on block-level U.S. Census data on 3-9 year-olds from 1990, 2000 and 2010 and supplemented by ESRI Sourcebook (24).

4 Charter schools are independently run and can adopt alternative pedagogical orientations even though they are publicly funded. Admissions to charter schools are not constrained by residence in a particular school district.

5 Isolation index at the school board level is used for the eight school boards of the Los Angeles Unified School District due to its large size.

6 Individual-level data on PBEs are unavailable due to privacy concerns. Expected count of PBEs (u_ij_) is calculated as: log(u_ij_) = log(t_ij_) + α + βx_ij_ + γ_zj_ + *u*_ij_, where schools are indexed by i and year by j, log(t_ij_) is the logged number of enrolled students (exposure), α is the global intercept, x_ij_ is a vector of lagged independent covariates measured by school and year with β estimated covariate effects, z is a set of indicator variables for year with γ estimated effects, and *u*ij is the error term, composed of *v*_i_~N(0,σV^2^) and ε_ij_~N(0,σ^2^).

7 Using different time lags, or no lag at all, does not substantially change the results (see Table A1 in Appendix for results using a 5-year lag; results from other models are available upon request).

8 Private schools are not included due to lack of detailed enrollment data by grade.

9 This finding may seem surprising as it appears to contradict traditional explanations linking lack of access to healthcare as a primary factor in under-vaccination.

**Table A1.** Correlates of PBEs among public school kindergartens, 5-year lag, 1998-2014^1^.

|  | Unadjusted<br>IRR<sup>2,4</sup> | Sig. | 95% C.I. |  | Adjusted<br>IRR<sup>4</sup> | Sig. | 95% C.I. |  |
| --- | --- | --- | --- | --- | --- | --- | --- | --- |
| Mother's education | 1.535*** | 0.000 | 1.508 | 1.563 | 1.186*** | 0.000 | 1.159 | 1.214 |
| Property values (ln) | 1.673*** | 0.000 | 1.596 | 1.753 | 0.989 | 0.670 | 0.940 | 1.040 |
| Pop. density (ln) | 0.669*** | 0.000 | 0.655 | 0.684 | 0.776*** | 0.000 | 0.759 | 0.792 |
| % white<sup>3</sup> | 1.344*** | 0.000 | 1.334 | 1.354 | 1.221*** | 0.000 | 1.208 | 1.233 |
| Physicians | 1.009*** | 0.000 | 1.007 | 1.011 | 1.004*** | 0.001 | 1.002 | 1.006 |
| Acupuncturists | 1.036*** | 0.000 | 1.029 | 1.043 | 0.998 | 0.509 | 0.991 | 1.005 |
| Chiropractors | 1.040*** | 0.000 | 1.035 | 1.045 | 1.026*** | 0.000 | 1.021 | 1.032 |
| Autism Diagnoses | 1.010*** | 0.000 | 1.004 | 1.016 | 1.004 | 0.184 | 0.998 | 1.010 |

Note:
<sup>1</sup> 2-level random intercept negative binomial regression model, N= 89,550 school-years. Year indicator variables included but not shown. All independent variables lagged 5 years.
<sup>2</sup> Adjusted for year indicator variables.
<sup>3</sup> Measured in 10% increments (deciles)
<sup>4</sup> Incidence risk ratio
†p < .10; \*p < .05; \*\*p < .01; \*\*\*p < .001

**Table A2.** Three-level model for correlates of PBEs among public school kindergartens, 2006-2014^1^.

|  | IRR<sup>3</sup> | Sig. | 95% C.I. |  |
| --- | --- | --- | --- | --- |
| Mother's education | 1.257*** | 0.000 | 1.222 | 1.293 |
| Property values (ln) | 1.294*** | 0.000 | 1.186 | 1.411 |
| Population density (ln) | 0.910*** | 0.000 | 0.884 | 0.937 |
| % white<sup>2</sup> | 1.158*** | 0.000 | 1.143 | 1.174 |
| Physicians | 0.999 | 0.166 | 0.997 | 1.001 |
| Acupuncturists | 0.970*** | 0.000 | 0.962 | 0.978 |
| Chiropractors | 1.030*** | 0.000 | 1.025 | 1.036 |
| Autism diagnosis | 1.007* | 0.025 | 1.001 | 1.012 |
| Autism advocacy orgs (county) | 0.996 | 0.105 | 0.992 | 1.001 |
| Pertussis rate, ages 0-6 (county) | 0.962† | 0.086 | 0.921 | 1.006 |

Note:
<sup>1</sup> 3-level random intercept Poisson model, N= 49,153 school-years. Year indicator variables included but not shown. All independent variables lagged 6 years.
<sup>2</sup> Measured in 10% increments (deciles)
<sup>3</sup> Incidence risk ratio
†p < .10; \*p < .05; \*\*p < .01; \*\*\*p < .001

**Table A3.** Effects of school type on PBEs among kindergartens, 1998-2014^1^Geometric overlap between EB tiles and wedges.

|  | IRR<sup>4</sup> | Sig. | 95% C.I. |  | IRR<sup>4</sup> | Sig. | 95% C.I. |  |
| --- | --- | --- | --- | --- | --- | --- | --- | --- |
| School type<sup>2</sup> |  |  |  |  |  |  |  |  |
| Private schools |  |  |  |  | 2.174*** | 0.000 | 2.074 | 2.279 |
| Charter schools |  |  |  |  | 3.347*** | 0.000 | 3.109 | 3.604 |
| Mother's education | 1.144*** | 0.000 | 1.119 | 1.170 | 1.145*** | 0.000 | 1.121 | 1.170 |
| Property value (ln) | 1.081** | 0.002 | 1.030 | 1.136 | 1.015 | 0.534 | 0.969 | 1.063 |
| Pop. density (ln) | 0.829*** | 0.000 | 0.812 | 0.846 | 0.807*** | 0.000 | 0.791 | 0.823 |
| % White<sup>3</sup> | 1.211*** | 0.000 | 1.200 | 1.223 | 1.204*** | 0.000 | 1.193 | 1.216 |

Note:
<sup>1</sup> 2-level random intercept negative binomial regression model, N= 116,928 school-years. Year indicator variables included but not shown. Demographic variables lagged 6 years.
<sup>2</sup> Reference category: public, non-charter schools.
<sup>3</sup> Measured in 10% increments (deciles)
<sup>4</sup> Incidence risk ratio
†p < .10; \*p < .05; \*\*p < .01; \*\*\*p < .001

**Table A4.**
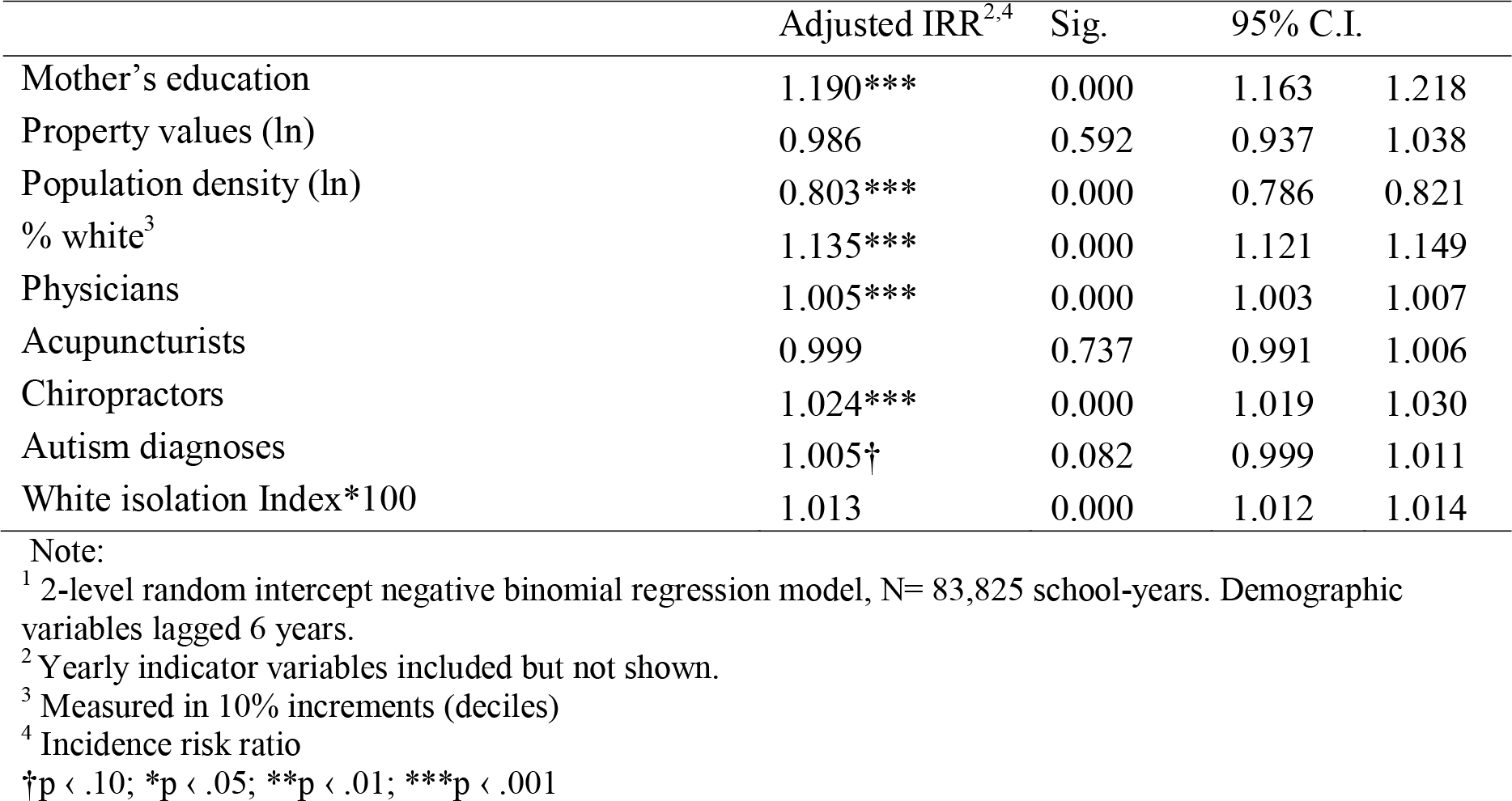
Effect of racial isolation on PBEs, rates among public kindergartens^1^.

|  | Adjusted IRR<sup>2,4</sup> | Sig. | 95% C.I. |  |
| --- | --- | --- | --- | --- |
| Mother's education | 1.190*** | 0.000 | 1.163 | 1.218 |
| Property values (ln) | 0.986 | 0.592 | 0.937 | 1.038 |
| Population density (ln) | 0.803*** | 0.000 | 0.786 | 0.821 |
| % white<sup>3</sup> | 1.135*** | 0.000 | 1.121 | 1.149 |
| Physicians | 1.005*** | 0.000 | 1.003 | 1.007 |
| Acupuncturists | 0.999 | 0.737 | 0.991 | 1.006 |
| Chiropractors | 1.024*** | 0.000 | 1.019 | 1.030 |
| Autism diagnoses | 1.005† | 0.082 | 0.999 | 1.011 |
| White isolation Index*100 | 1.013 | 0.000 | 1.012 | 1.014 |

Note:
<sup>1</sup> 2-level random intercept negative binomial regression model, N= 83,825 school-years. Demographic variables lagged 6 years.
<sup>2</sup> Yearly indicator variables included but not shown.
<sup>3</sup> Measured in 10% increments (deciles)
<sup>4</sup> Incidence risk ratio
†p < .10; \*p < .05; \*\*p < .01; \*\*\*p < .001

**Table A5.**
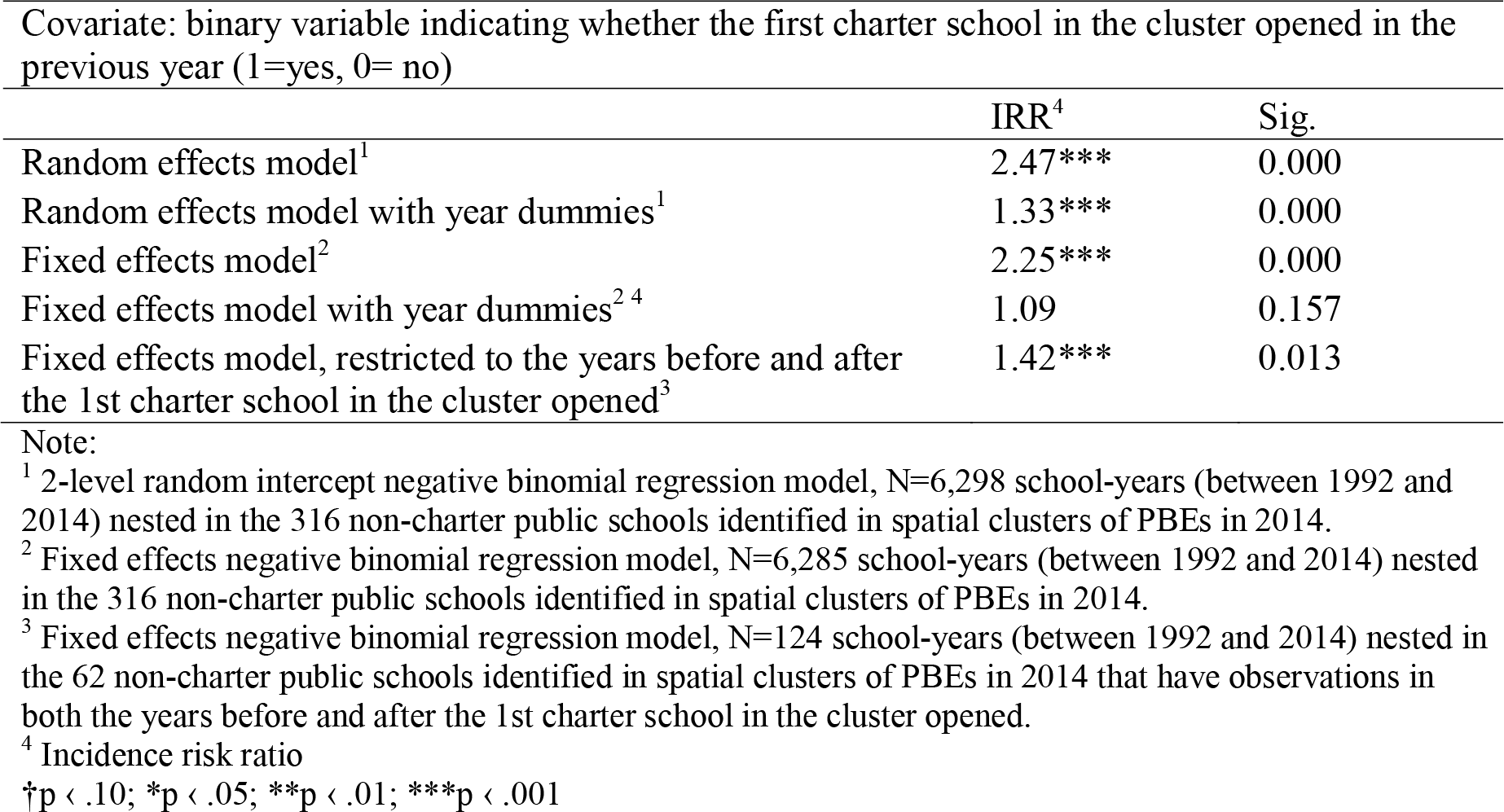
Fixed and random effects models of panel data of public non-charter schools in 2014 clusters.

**Figure A1.**
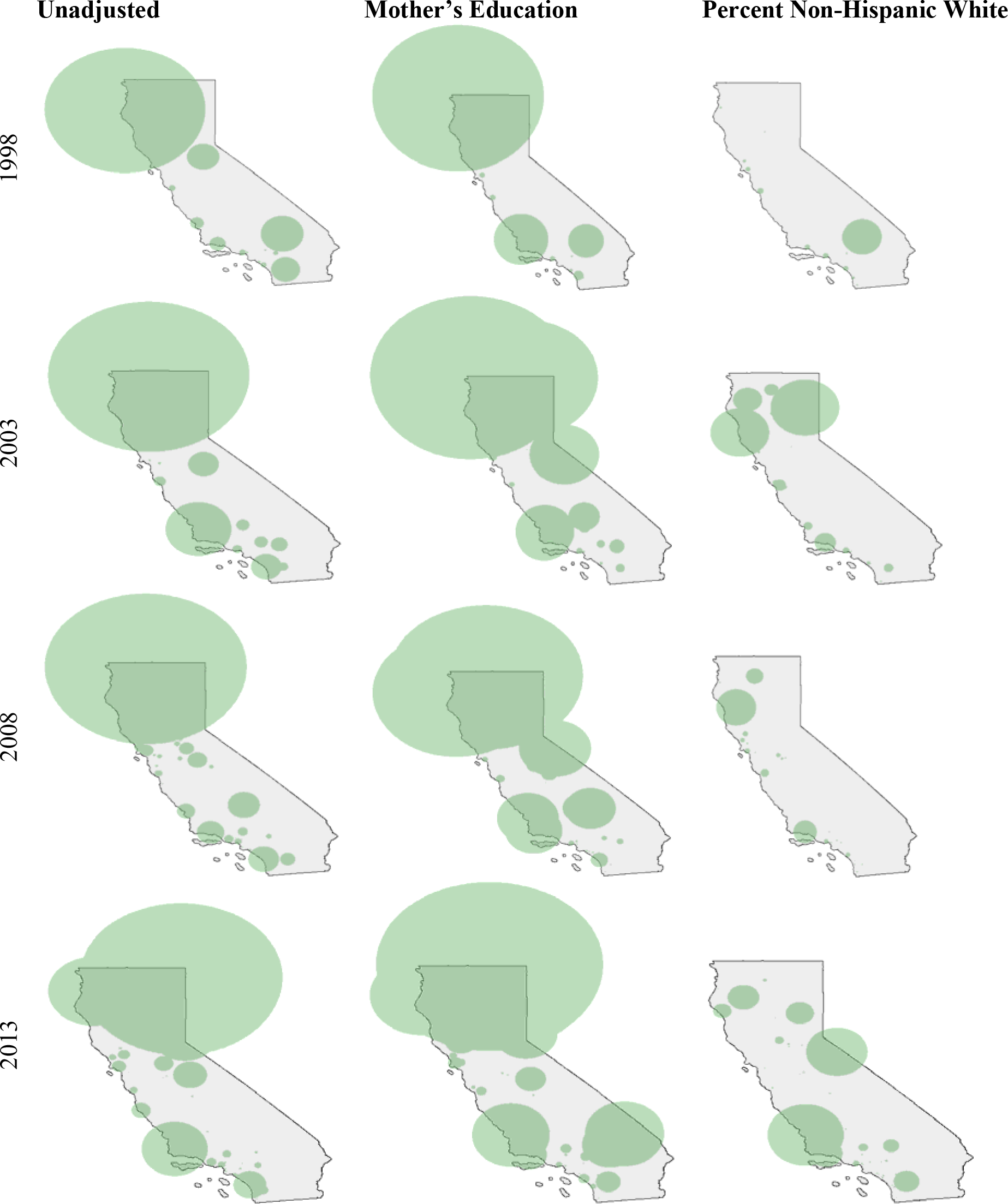
Adjusted spatial clusters of PBEs by Mother’s Education and Percent Non-Hispanic White, 1998-2013.

**Figure A2.**
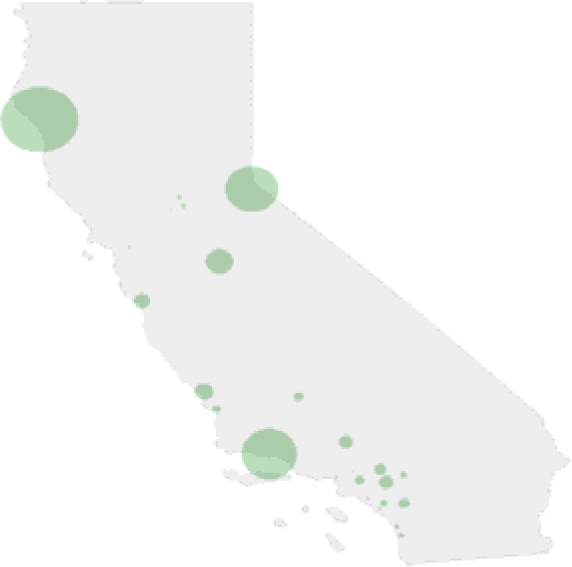
Spatial clusters of PBEs across public schools adjusted for all covariates, with charter schools removed, 2014.

## References

1 Dubé E, et al. (2013) Vaccine hesitancy. Human vaccines & immunotherapeutics 9(8):1763–1773.

2 Sadaf, A, Richards, JL, Gians J, Salmon, DA, & Omer SB (2013) A systematic review of interventions for reducing parental vaccine refusal and vaccine hesitancy. Vaccine 31(40):4293–4304.

3 Constable, C, Blank, NR, & Caplan, AL (2014) Rising rates of vaccine exemptions: problems with current policy and more promising remedies. Vaccine 32(16):1793–1797.

4 Eames, KT (2009) Networks of influence and infection: parental choices and childhood disease. journal of the Koyal Society, Interface / the Royal Society 6(38): 811–814.

5 Salathe, M & Bonhoeffer, S (2008) The effect of opinion clustering on disease outbreaks. journal of the Royal Society, Interface / the Royal Society 5(29):1505–1508.

6 Masumdar, S, King, M, Liu K-Y, Zerubavel, N, & Bearman, P (2010) The spatial structure of autism in California, 1993-2001. Health & Place 16(3):539–546.

7 Masumdar, S, Winter, A, Liu, KY, & Bearman, P (2013) Spatial clusters of autism births and diagnoses point to contextual drivers of increased prevalence. Soc Sri Med 95:87–96.

8 Liu, K, King, M, & Bearman, PS (2010) Social Influence and the Autism Epidemic. American journal of Sociology 115(5):1387–1434.

9 Fountain C & Bearman PS (2011) Risk as Social Context: Immigration Policy and Autism in California. Sociologicall Forum 26(2 %L 0000).

10 Taylor, LE, Swerdfeger AL, & Eslick, GD (2014) Vaccines are not associated with autism: an evidence-based meta-analysis of case-control and cohort studies. Vaccine 32(29):3623–3629.

11 Freed, GL, Clark, SJ, Butchart, AT, Singer, DC, & Davis, MM (2010) Parental Vaccine Safety Concerns in 2009. Pediatrics 125(4):654–659.

12 Stefanoff, P, et al. (2010) Tracking parental attitudes on vaccination across European countries: The Vaccine Safety, Attitudes, Training and Communication Project (VACSATC). Vaccine 28(35):5731–5737.

13 Schwarser, R (1994) Optimism, Vulnerability, and self-beliefs as health-related cognitions: A systematic overview. Psychohsy & Health 9(3):161–180.

14 Safi H, et al. (2012) Vaccine policy and Arkansas childhood immunisation exemptions: a multi-year review. Am J Prev Med 42(6):602–605.

15 Omer, SB, et al. (2008) Geographic clustering of nonmedical exemptions to school immunisation requirements and associations with geographic clustering of pertussis. Am J Epidemiol 168(12):1389–1396.

16 Atwell JE, et al. (2013) Nonmedical vaccine exemptions and pertussis in California, 2010. Pediatrics 132(4):624–630.

17 Imdad, A, et al. (2013) Religious exemptions for immunisation and risk of pertussis in New^7^ York State, 2000-2011. Pediatrics 132(1)37–43.

18 Carrel, M & Bitterman, P (2015) Personal Belief Exemptions to Vaccination in California: A Spatial Analysis. Pediatrics.

19 Lieu, TA, Ray, GT, Klein, NP, Chung, C, & Kulldorff, M (2015) Geographic Clusters in Underimmunisation and Vaccine Refusal. Pediatrics.

20 Richards JL, et al. (2013) Nonmedical exemptions to immunisation requirements in California: A 16-year longitudinal analysis of trends and associated community factors. Vaccine 31(29):3009–3013.

21 Birnbaum, MS, Jacobs, ET, Ralston-King J, & Ernst, KC (2013) Correlates of high vaccination exemption rates among kindergartens. Vaccine 31(5):750–756.

22 Buttenheim, AM, et al (2015) MMR vaccination status of children exempted from school entry immunisation mandates. Vaccine 33(46):6250–6256.

23 Croen, LA, Grether JK, Hoogstrate J, & Selvin, S (2002) The changing prevalence of autism in California. J Autism Dev Disord 32(3):207–215.

24 ESRI (2002-2004) Community Sourcebook - America with ArcReader (2002-2004 editions). (ESRI, Redlands, CA).

25 Bauch, CT & Bhattacharyya, S (2012) Evolutionary game theory and social learning can determine how vaccine scares unfold. PLOS computational biology 8(4): e1002452.

26 Brennan, JM, et al (2017) Trends in Personal Belief Exemption Rates Among Alternative Private Schools: Waldorf, Montessori, and Holistic Kindergartens in California, 2000-2014. Am J Public Health 107(1):108–112.

27 Salmon, DA. et al (2005) Factors Associated With Refusal of Childhood Vaccines Among Parents of School-aged Children: A Case-Control Study. Arch Pediatr Adoksc Med 159(5):470–476.

28 Walker ET & Rea CM (2016) Pediatric Care Provider Density and Personal Belief Exemptions From Vaccine Requirements in California Kindergartens. American journal of Public Health 106(7):1336–1341.

29 Campbell, JB, Busse, JW, & Injeyan HS (2000) Chiropractors and vaccination: A historical perspective. Pediatrics 105(4):E43.

30 Brown KF, et al (2010) Factors underlying parental decisions about combination childhood vaccinations including MMR: A systematic review. Vaccine 28(26):4235–4248.

31 Shui, IM, Weintraub ES, & Gust, DA (2006) Parents Concerned About Vaccine Safety: Differences in Race/Ethnicity and Attitudes. American journal of Preventive Medicine 31(3):244–251.

32 Massey, DS & Denton, NA (1988) The Dimensions of Residential Segregation. Social Forces 67(2)581–315.

33 Kulldorff, M (1999) Spatial Scan Statistics: Models,Calculations, and Applications. Scan Statistics and Applications, Statistics for Industry and Technology, eds Glas J & Balakrishnan N (Birkhäuser Boston), pp 303–322.

34 Buttenheim, A, Jones M, & Baras Y (2012) Exposure of California kindergartners to students with personal belief exemptions from mandated school entry vaccinations. Am J Public Health 102 (8):e59–67.

35 Ilinman, A (1999) Eradication of vaccine-preventable diseases. Annu Rev Public Health 20:211–229.

36 Richards, JL, et al (2013) Nonmedical exemptions to immunisation requirements in California: a 16-year longitudinal analysis of trends and associated community factors. Vaccine 31(29)3009–3013.

37 Yang TY & Silverman R (2015) Legislative Prescriptions for Controlling Nonmedical Vaccine Exemptions. journal of the American Medical Association 313(3):247–248.

38 Delamater, PL, Leslie, TF, & Yang, Y (2017) Change in medical exemptions from immunisation in California after elimination of personal belief exemptions. JAMA 318(9):863–864.

39 Seipel, T (2016/06/30) California’s vaccine law: Opponents moving, home schooling to avoid controversial mandate. Mercury News.

40 Gumbel, A (2015/01/25/) Disneyland measles outbreak leaves many anti-vaccination parents unmoved. The Guardian, Section US News.

41 Eyal, G (2010) The autism matrix: the social origins of the autism epidemic (Polity, Cambridge; Malden, MA) pp viii, 312 p.

42 Gaudino, JA & Robison, S (2012) Risk factors associated with parents claiming personal-belief exemptions to school immunisation requirements: community and other influences on more skeptical parents in Oregon, 2006. Vaccine 30(6):1132–1142.

43 Prislin, R, Dyer, JA, Blakely CH, & Johnson CD (1998) Immunisation status and sociodemographic characteristics: the mediating role of beliefs, attitudes, and perceived control. Am J Public Health 88(12):1821–1826.

44 Centola, D (2010) The Spread of Behavior in an Online Social Network Experiment. Science 329(5996):1194–1197.

45 Granovetter, M (1978) Threshold Models of Collective Behavior. American journal of Sociology 83(6):1420–1443.

46 McPherson, M, Smith-Lovin L, & Cook, JM (2001) Birds of a feather: llomophily in social networks. Annual Review of Sociology 27:415–444.

47 Centola, D & Macy, M (2007) Complex contagions and the weakness of long ties. American Journal of Sociology 113(3):702–734.

48 Horne, Z, Powell, D, Hummel, JE, & Holyoak, KJ (2015) Countering antivaccination attitudes. Proceedings of the National Academy of Sciences 112(33):10321–10324.

49 Valente TW (2012) Network interventions. Science 337(6090):49–53.

